# Unraveling the differential dynamics of developmental fate in central and peripheral nervous systems

**DOI:** 10.1101/072579

**Authors:** Dola Sengupta, Sandip Kar

## Abstract

Bone morphogenetic protein 2 (BMP2), differentially regulates the developmental lineage commitment of neural stem cells (NSC’s) in central and peripheral nervous systems. However, the precise mechanism beneath such observations still remains illusive. To decipher the intricacies of this mechanism, we propose a generic mathematical model of BMP2 driven differentiation regulation of NSC’s. The model efficiently captures the dynamics of the wild-type as well as various mutant and over-expression phenotypes for NSC’s in central nervous system. Our model predicts that the differential developmental dynamics of the NSC’s in peripheral nervous system can be reconciled by altering the relative positions of the two mutually interconnected bi-unstable switches inherently present in the steady state dynamics of the crucial developmental fate regulatory proteins as a function of BMP2 dose. This model thus provides a novel mechanistic insight and has the potential to deliver exciting therapeutic strategies for neuronal regeneration from NSC’s of different origin.

---

Bone morphogenetic protein 2 (BMP2), a pleotropic cytokine, is crucial to determine the developmental fates of neural progenitor cells in central and peripheral nervous systems. Telencephalic neural progenitor cells acquired from central nervous system (CNS) switch from neuronal to gliogenic developmental fate under transient exposure of BMP2^1,2^. On the contrary, neural crest stem cells obtained from peripheral nervous system (PNS) differentiate into neurons with increased doses of BMP2 in the cell culture medium^1,3–5^. Although most of the transcription factors regulating these developmental events in neural stem cells (NSC’s) are well characterized in literature^1,2,4,6–8^, the precise dynamics that governs the BMP2 driven developmental fate in CNS and PNS still remained as an unsolved puzzle. To translate this kind of basic biological knowledge into a therapeutic application such as neuronal regeneration after nerve injuries in brain and spinal cord by employing a certain dose of BMP2 in a context dependent manner, it is imperative to have a detailed understanding of how BMP2 regulates the differentiation of the neural stem cells into neurons in CNS and PNS.

Undoubtedly, the underlying gene regulatory network organizing the developmental event in the nervous system is extremely complex. Activities of various basic helix– loop–helix (bHLH) transcription factors such as Mash1 (mammalian achaete-scute homologue) and Ngn (neurogenin) control the lineage specific neuronal cell fate in CNS and PNS^1–4,9^. Another set of HLH factors Hes1 (homologue of hairy and Enhancer of Split) and Hes5 negatively regulate activities of these bHLH proteins^1,6,10,11^. These negative HLH proteins along with Id1 (inhibitor of differentiation) protein contribute to the antineurogenic effect of BMP2 by reinforcing gliogenesis^1,2^ whereas proneural bHLH proteins Mash1 and Ngn are the master regulators of neuronal differentiation. Surprisingly, another member of Hes family of proteins, Hes6, has antagonistic effect on Hes1 and Hes5^6,7,9,12^. In this context, it is important to note that more than one signal transduction pathways modulate these transcriptional regulators in different extent to orchestrate the overall developmental process in different cell types.

Keeping these complexities in mind, one can presume that mathematical and computational modeling approaches might provide some initial insight to disentangle the underlying regulatory dynamics of this complex network. No wonder, in last few decades, mathematical and computational modeling studies were used extensively to understand the dynamic regulation of few of these important transcription factors (such as Hes1) either in isolation, or taking a part of the gene regulatory network organizing the developmental events^13–15^ in NSC’s of different origin. As of now, none of the earlier studies investigated the differential role played by BMP2 in controlling the developmental fate in central and peripheral nervous systems. This article puts forward an extensive and unified mathematical model of BMP2 driven neuronal stem cell differentiation regulation that accounts for the differential nature of developmental dynamics of NSC’s in central and peripheral nervous systems.

We divide the entire gene regulatory network controlling the BMP2 driven differentiation of NSC’s in CNS and PNS into three simple gene interaction modules. By using extensive bifurcation analysis, we integrated the mathematical models of the individual modules one by one systematically to build up the overall model of BMP2 driven neural stem cell differentiation for CNS. An exhaustive list of mutant and overexpression studies available in the experimental literature are utilized to establish the mathematical model that qualitatively describes how BMP2 alters the cell fate decision during development of neural progenitor cells in CNS. To this end, we employ a systematic sensitivity analysis on the model parameters and predict the possible routes to obtain a PNS like behavior using the same model developed for CNS. Our model predicts several mutant and over-expression phenotypes for NSC’s acquired from PNS and importantly unveils more than one possibility to efficiently transform the NSC’s into neuron in a lineage specific manner. Existing experimental literature corroborates well with these model predictions^16–18^. We believe that this generic model provides an important preliminary insight and satisfactorily explains the differential regulation of developmental lineage commitment in CNS and PNS by BMP2 as a purely dynamical effect.

## The Model of BMP2 driven neuronal differentiation regulation

The model, depicted in Fig. 1 (detail description is provided in Supplementary Fig. 1, SI Text and *SI Appendix*), comprised of three individual modules that describes how BMP2 controls the process of development in the NSC’s of CNS and PNS. BMP2 is known to regulate the transcription of Mash1 and Id1 proteins through direct and indirect mechanisms^1,2,4^ as shown in module-1 wherein Mash1 can further activate its own transcription with the help of E47 protein^1,19^. Id1 on the other hand sequesters E47 through competitive binding and can inhibit Mash1 by actively degrading Mash1^2^. In module-2, BMP2 influences only the transcription of Hes5 protein^1^, which is also known to regulate its own transcription through a negative feedback mechanism similar to Hes1 protein^8,20–24^. Furthermore, Hes5 has a mutually antagonistic relationship with Ngn protein responsible for neurogenesis^6^. In addition, Ngn activates the Hes6 protein, which in turn initiates the degradation process of Hes5^6,7,12,25,26^. Module-3 is rather simple and only described by a negative feedback loop of Hes1 protein on its own transcription^14,27^. BMP2 does not have any direct connection with Hes1 protein activation^1^ but later we will see that this module-3 plays a crucial role in determining the oscillatory dynamics observed in this kind of system. The three modules are interconnected through multiple regulatory interactions, which are further described in details in SI Text and *SI Appendix*. There we have more comprehensive schemes for modules 1-3 in *SI Appendix* with detail description of each module in terms of precise molecular interaction curated from available experimental literature.

**Figure 1.**
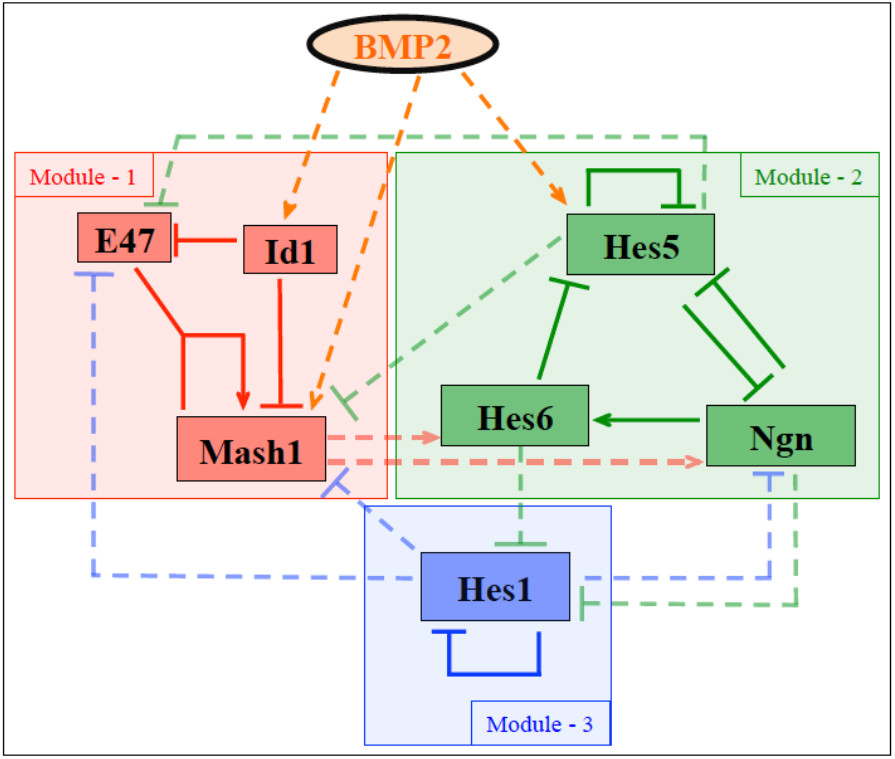
Simplified scheme of the neuronal differentiation regulation network. (Arrows (Solid - intra modular, dashed – inter modular) and hammer-headed lines (Solid - intra modular, dashed – inter modular) represent direct or indirect activation and inhibition processes respectively.) The overall network is divided into three modules and involves neurogenic HLH transcription factors Mash1, Ngn, positive HLH factor Hes6 along with negative HLH factors Id1, Hes5 and Hes1. The effect of BMP2 on Mash1, Id1 and Hes5 is incorporated in module-1 and module-2. The detailed regulatory interactions are described in *SI Appendix* where comprehensive schemes for the whole interaction network (Supplementary Fig. 1) and three modules are discussed elaborately. Corresponding kinetic equations, description of the variables and parameters are depicted in SI Text and *SI Appendix*.

The overall model (Fig. 1, Supplementary Fig. 1) can be represented by ordinary differential equations based on mass action kinetic terms (with only two Michealis-Menten terms used in module–2 and 3) as provided in SI Text (comprising of 46 kinetic equations). All the variables, the definition of the parameters and their numerical values are provided in SI Text. In this regard, it is worthwhile mentioning that we gathered the values for several parameters from experimental papers^15,27–31^ and used reasonable guess values for the others to obtain the dynamic features of the concerned system. Initially, we focused more on establishing a mathematical model that will convincingly describe not only the WT features but also simulate most of the mutants and overexpression phenotypes for the NSC’s under different doses of BMP2 in CNS, as more experimental results are available for cells derived from CNS. Once we develop this mathematical model for CNS, the main aim of this study is to understand whether this same mathematical model (with no changes in the network architecture or interactions) with or without any significant changes in the model parameters (as few in numbers as possible), can manifest a PNS like behavior for increasing doses of BMP2 or not?

## Results

### Increasing levels of BMP2 switches the developmental fate of WT NSC’s in CNS

To understand the characteristic dynamical transitions under the influence of BMP2, we first perform the bifurcation analysis (Fig. 2a Supplementary Fig. 2a) for the total model (given in SI Text) that describes how the total Mash1 and Hes5 protein concentrations are affected by increasing BMP2 dose. In Fig. 2a (left panel, Mash1 and right panel, Hes5) and Supplementary Fig. 2a, it is evident that the steady state levels of total Mash1 and Hes5 proteins as a function of BMP2 still contain two interconnected switches (as seen earlier for module–1, see *SI Appendix* for details) but now in the total model both the lower and upper steady states are unstable. The saddle node SN_1_ in Fig. 2a appears for very low dose of BMP2 (∼ 0.186 s.u.). At BMP2=0.827 s.u., the system undergoes a Hopf bifurcation (HB_2_) that makes the dynamics beyond this value of BMP2 oscillatory. Although BMP2 controls the transcription of *Mash1*, *Id1* and *Hes5* genes, initially when BMP2 level is very low it affects the *Mash1* transcription by up regulating the E47 mediated *Mash1* transcription more than the *Id1* and *Hes5* transcriptions.

**Figure 2.**
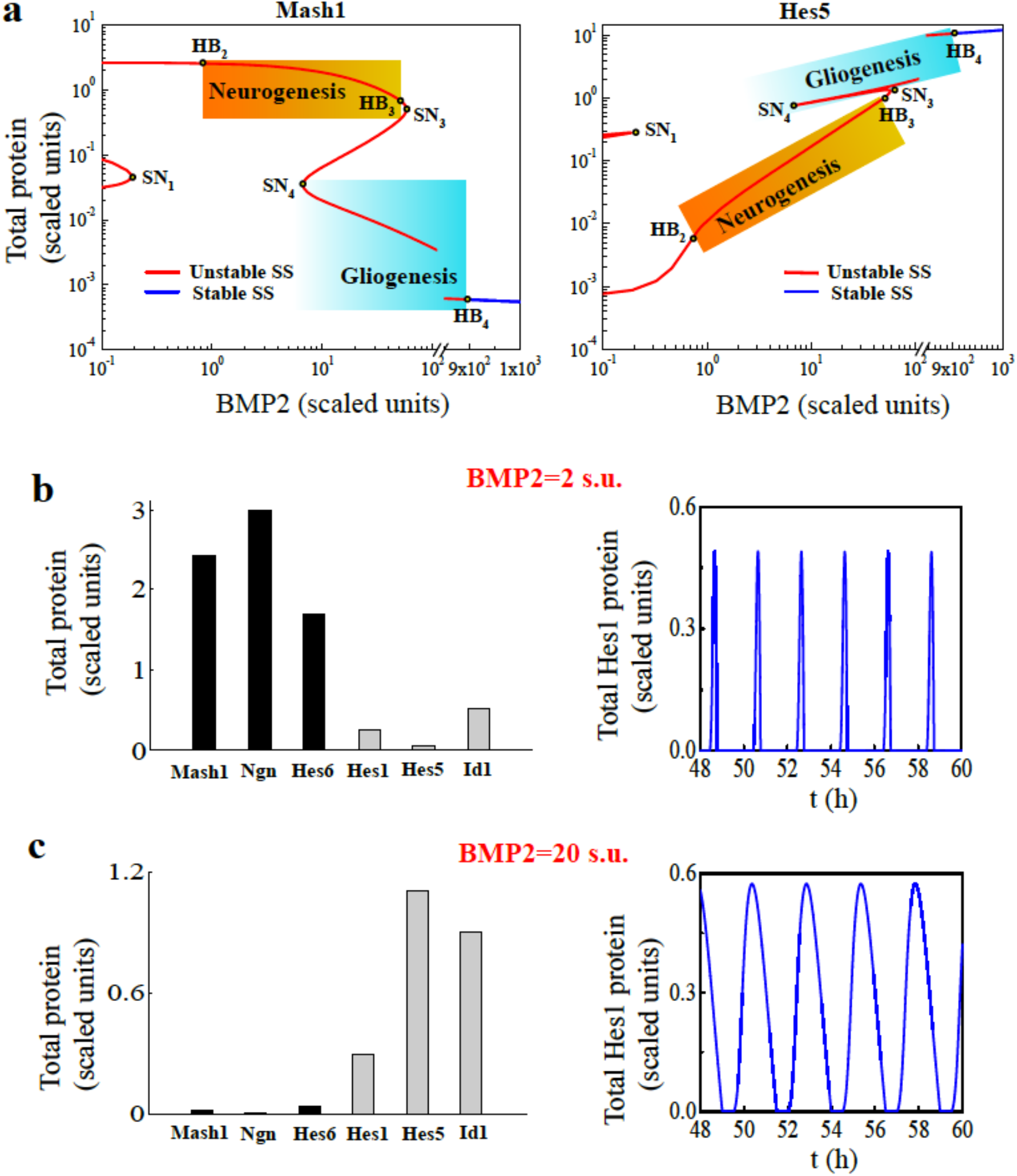
Bmp2 driven developmental lineage commitment of WT NSC’s in CNS. **(a)** Bifurcation diagrams of total Mash1 and Hes5 proteins are plotted as a function of BMP2 (both the axes are shown in log scale). Increasing the level of BMP2 drives the developmental cell fate from neurogenic state (high Mash1 and low Hes5 expressions, shown by the graded orange region) to gliogenic state (low Mash1 and high Hes5 expressions, shown by the graded blue region). **(b)** Expression levels of the proneural (Mash1, Ngn and Hes6), gliogenic (Hes5 and Id1) and Hes1 proteins at BMP2=2 s.u. show clear indication of neuronal fate commitment (left panel) and Hes1 protein concentration shows ∼2 h oscillation period (right panel). **(c)** Expression levels of the proneural (Mash1, Ngn and Hes6), gliogenic (Hes5 and Id1) and Hes1 proteins at BMP2=20 s.u. evidently indicate gliogenic fate commitment (left panel) and Hes1 protein concentration shows ∼2 h oscillation period (right panel). Since all the proteins are showing oscillatory behavior for both low and high doses of BMP2, we plotted the expression levels of the individual proteins by taking the average of the oscillation amplitude. The observations were made on day 2 to corroborate with the experimental procedures. The parameter values are given in SI Text.

Thus, for low values of BMP2 the system quickly attains high levels of Total Mash1 protein (neuronal state). Once there is enough accumulation of Id1 and Hes5 proteins in the system, they start to affect the Mash1 level by initiating its degradation (Id1 mediated) as well as sequester the E47 protein (both Id1, Hes5 and even Hes1) to inhibit the Mash1 positive feedback to its own transcription. This will make sure that the Mash1 level starts dropping after BMP2=0.827 s.u. and cause the 2^nd^ switch in the process with saddle nodes SN_3_ (BMP2 ∼ 58.14 s.u.) and SN_4_ (BMP2 ∼ 6.71 s.u.) (Fig. 2a, Supplementary Fig. 2a). The lower (for Mash1) and upper steady states (for Hes5) starting at BMP2 ∼ 6.71 s.u. in Fig. 2a are also unstable and give rise to stable oscillations for further higher values of BMP2 due to presence of Hopf bifurcation (HB_4_) at BMP2 ∼ 895.7 s.u..

At this point, we define BMP2=2 s.u. as one of the representative low dose of BMP2 and BMP2=20 s.u. as high dose of BMP2 to observe the expression levels of different proteins in the network under high and low doses of BMP2. Although this definition seems arbitrary to begin with, it is a reasonable one and we stick to this definition through out our study as in the experimental papers the high dose of BMP2 (∼80 ng/ml BMP2 dose used in experiments) is considered to be at least 10 times higher than the low dose^1,3,4^. Now if we initialize the dynamical system at very low dose (close to 0) of BMP2 (representative of cells in culture medium with almost no BMP2) and suddenly increase the level of BMP2 to BMP2=2 s.u., we obtain the expression levels of the different proteins as shown in Fig. 2b (left panel). High levels of pro-neural (Mash1, Ngn and Hes6) proteins and low levels of gliogenic (Hes5 and Id1) proteins (Fig. 2b, left panel) definitely indicate towards the neuronal phenotype under lower dose of BMP2 (2 s.u.). Importantly the Hes1 protein concentration seems to oscillate with a period of ∼2 h (Fig. 2b, right panel) which successfully corroborate with the experimental observations^27,32^.

On the contrary, if we initialize the dynamical system similarly (for BMP2 close 0) and suddenly employ BMP2=20 s.u., the system preferably leans towards a gliogenic phenotype indicated by low levels of pro-neural proteins and higher levels of gliogenic protein (Fig. 2c, left panel) maintaining the Hes1 oscillation (Fig. 2c, right panel) with a period of ∼2 h intact. Interestingly, the Hes1 oscillation amplitude under low and high doses of BMP2 remains similar as observed in experiments^1^. This situation clearly resembles the way BMP2 drives developmental fate of WT neuronal stem cells obtained from CNS in experiments^1,2^. Our model further predicts that the phenotypic outcome for sudden changes in the BMP2 concentration (Supplementary Fig. 2b) after keeping the cells in a very low dose of BMP2 (Close to 0) initially will be quite different if the BMP2 concentration in the cell culture medium is increased in a continuous fashion (Supplementary Fig. 2c). Consequently, in the later case, the transition of NSC’s from neuronal to gliogenic fate will happen at much higher BMP2 level (Supplementary Fig. 2c). One needs to verify this prediction experimentally as such kind of experiments has not been performed yet in literature.

### The model describes a number of experimental phenotypes for NSC’s in CNS

The model developed so far is capable of capturing a number of phenotypic situations observed in different experiments performed with NCS’s obtained from CNS under different BMP2 conditions. Different experimental conditions and the corresponding observed phenotypes are listed in Table –1 along with the list of figures showing the successfully reproduced model simulated phenotypes. It has been observed experimentally that single serum treatment instigates *Hes1* mRNA oscillation with 2-3 hour cycle in various cultured cells like fibroblasts (C3H10T1/2), myoblasts (C2C12), neuroblastoma cells (PC12) and teratocarcinoma cells (F9) and there exists a time delay of the order of 15-30 minutes between Hes1 protein and *Hes1* mRNA oscillations^27^. The model simulations with BMP2=20 s.u. (as well as with 2 s.u.) evidently show 2 h period oscillation of *Hes1* mRNA (Supplementary Fig. 3a) and protein (Fig. 2b-c) with a time delay of ∼25 minutes (Fig. 3a). In this context, it is important to mention that previous attempts to model Hes1 oscillations had mostly used an explicit time lag (τ) in the differential equations of Hes1 protein and mRNA to account for this kind of time delay^13–15^.

**Figure 3.**
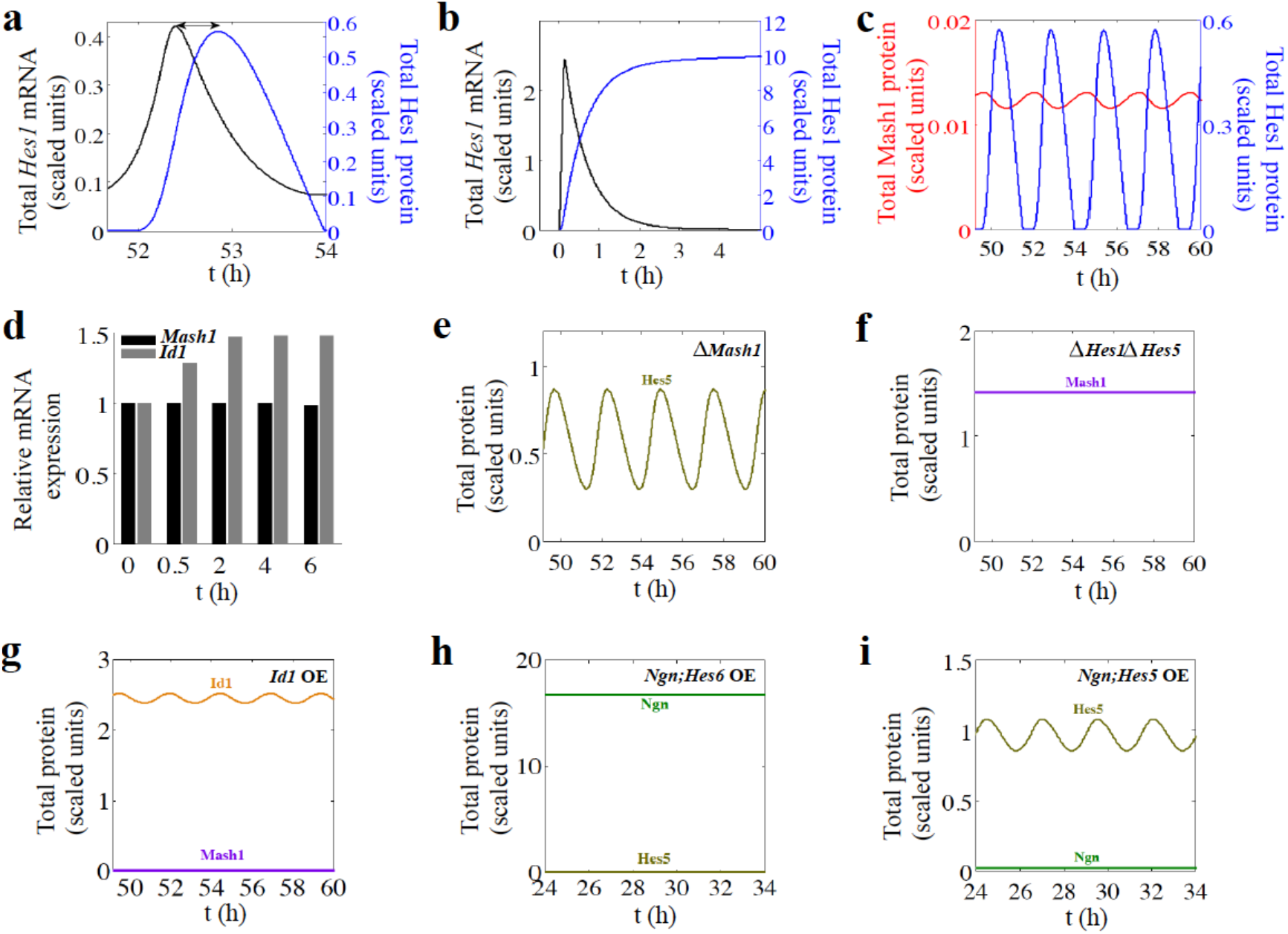
Model simulates a number of experimentally observed phenotypes in CNS. **(a)** Total Hes1 protein oscillation is delayed relative to *Hes1* mRNA oscillation by ∼25 minutes and peak to trough ratio is ∼2.3. **(b)** Degradation of Hes1 protein is stopped (*k*_dhes1_=*k*_dyn_=0 min^-1^) to incorporate the effect of the presence of proteasome inhibitor. This results in constant repression of *Hes1* transcription by highly expressing Hes1 protein level. **(c)** Mash1 protein expression oscillates in the opposite phase to Hes1. (a-c) Simulations were done at BMP2=20 s.u. **(d)** Relative expression levels of *Id1* and *Mash1* mRNA’s. BMP2 induces increase in *Id1* mRNA level (at BMP2=10 s.u. with respect to basal level, BMP2=0.5 s.u.) whereas *Mash1* mRNA does not show any significant difference up to 6 h of BMP2 addition under similar condition. **(e)** Deletion of *Mash1* gene, *G*_mt_=0 s.u., results in the up-regulation of Hes5, which indicates inhibition of neurogenesis even at low dose of BMP2 (BMP2=2 s.u.). **(f)** *Hes1*;*Hes5* double knockout (*G*_t_=*G*_5t_=0 s.u.) elevates Mash1 level even at high dose of BMP2 (BMP2=20 s.u.) leading to neurogenesis. **(g)** Overexpression of *Id1* gene (*G*_idt_=5 X WT) at low BMP2 (BMP2=2 s.u.) down-regulates Mash1 level resulting in gliogenesis. **(h)** Simultaneous overexpression of *Ngn* and *Hes6* genes (*G*_nt_=*G*_ht_=5 X WT) causes neurogenesis even at high dose of BMP2 (BMP2=20 s.u.) with decrease in Hes5 level. **(i)** Simultaneous overexpression of *Ngn* and *Hes5* genes (*G*_nt_=*G*_5t_=5 X WT) predicts the gliogenic fate of the system at low dose of BMP2 (BMP2=2 s.u.) with increase in Hes5 level. Other parameters are same as mentioned in SI Text.

**Table-1.**
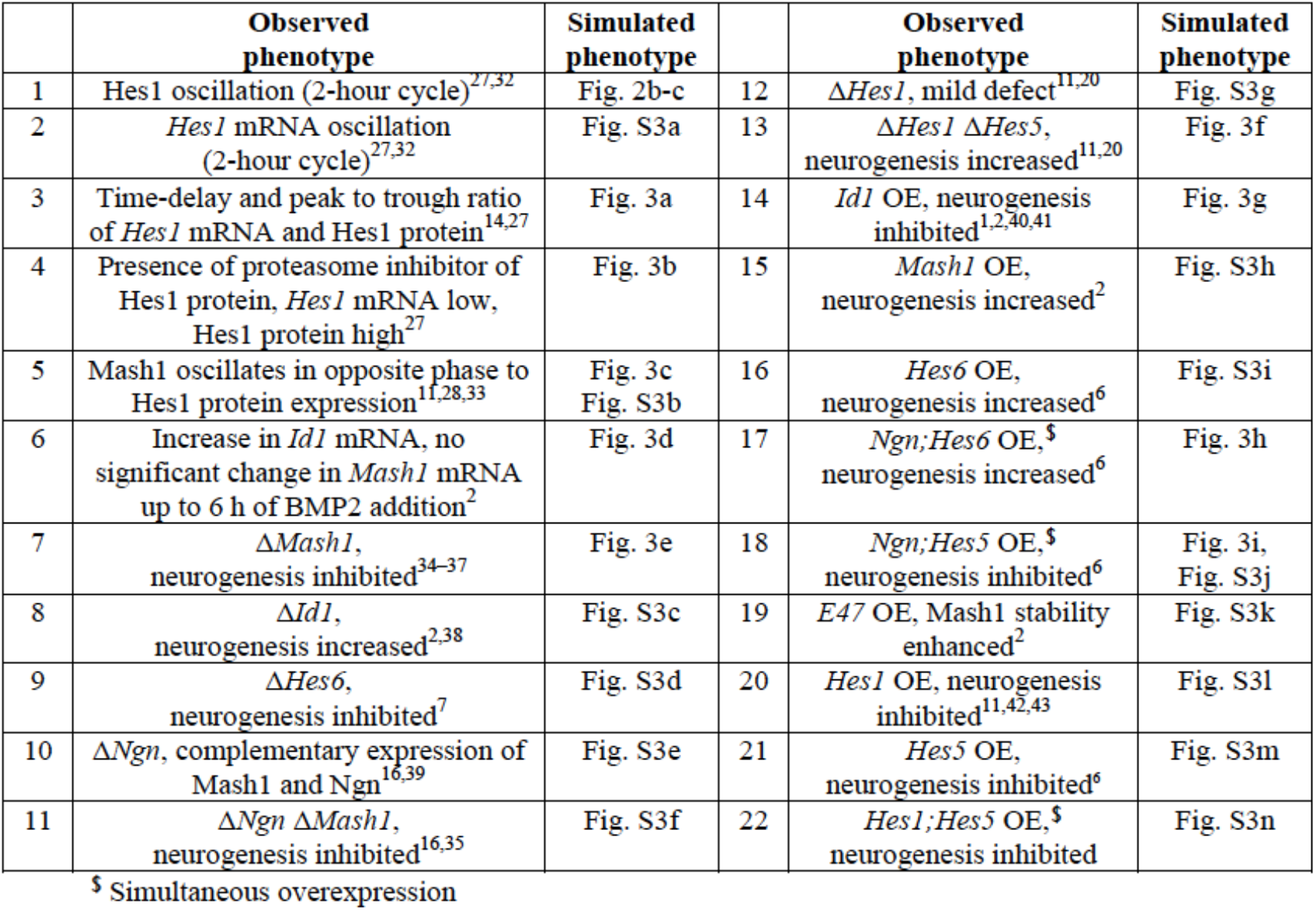
List of Mutant and over-expression phenotypes reproduced by the model.

In our model the time delay comes in automatically because an appropriate molecular mechanism (Supplementary Fig. 1) is used to model the negative feedback present in the Hes1 dynamics. The experimental peak to trough ratio^27^ (∼2.3) of the *Hes1* mRNA and protein can also be reproduced from the model simulation (Fig. 3a). Hirata et al. showed that inhibition of degradation of the Hes1 protein by using a proteasome inhibitor MG132 generates a transient increase in *Hes1* mRNA level at high serum dose but after a while it leads to continuous suppression of *Hes1* mRNA^27^. This observation can be nicely shown from our model simulation by stopping the degradation of Hes1 protein (Fig. 3b). Furthermore, our model simulation accounts for the experimental findings^28,33^ that the Mash1 protein level oscillates in the opposite phase of the Hes1 oscillation (Fig. 3c Supplementary Fig. 3b). In Fig. 3d, we show model simulated relative expression levels of *Id1* and *Mash1* mRNA’s, which corroborates with the experimental finding that BMP2 induces increase in *Id1* mRNA level (with respect to basal expression level) whereas *Mash1* mRNA does not show any significant difference up to 6 hours of BMP2 addition^2^.

### The model conforms to a number of mutant phenotypes for NSC’s in CNS

We have performed model simulations for different single and double mutant cases for the NSC’s derived from CNS and obtain excellent agreement with the experimental results. To begin with, we simulated the Δ*Mash1* mutant for low dose of BMP2 (Fig. 3e) and it caused an elevated level of Hes5 protein than the WT case (Fig. 2b, right panel) indicating inhibition of neurogenesis and onset of gliogenesis. This corroborates with the experimental fact that Δ*Mash1* NSC’s from CNS are more prone to gliogenic fate with increased expression level of Hes5 protein^34–37^. Similar to this observation, Jhas et al. had shown that neurogenesis is inhibited in Δ*Hes6* NSC’s originated from CNS^7^ and model simulation performed under such condition (Supplementary Fig. 3d) matches with the experimental finding. On the contrary, Δ*Id1* cells from CNS leads to premature neuronal differentiation^2,38^ and our model simulation (Supplementary Fig. 3c) can faithfully demonstrate this fact as well. Interestingly, it has been observed in experiments that Δ *Ngn* cells obtain from CNS still show reasonable level of Mash1 expression and consequently leans toward neuronal fate^16,39^. Model simulations performed under Δ*Ngn* condition (Supplementary Fig. 3e) retains the higher expression levels of Mash1 protein (in comparison to WT (Fig. 2b)) as seen in experiments^16,39^ (sign of neurogenesis) but model simulation under a double mutant condition Δ*Ngn* Δ*Mash1* (Supplementary Fig. 3f) predicts that the system will be pushed towards gliogenesis. In experiments neurogenesis is completely inhibited and gliogenesis is favored under Δ*Ngn* Δ*Mash1* mutant condition^16,35^. Ohtsuka et al. further demonstrated that Δ*Hes1* mutant has a mild defect and NSC’s from CNS were still adopting a gliogenic fate but a double mutant in the form of Δ*Hes1* Δ *Hes5* is lethal and causes increased level of neurogenesis^20^. The model simulation for Δ*Hes1* mutant (Supplementary Fig. 3g, with high level of Hes5 protein) and Δ*Hes1* Δ*Hes5* mutant (Fig. 3f, with high level of Mash1 protein) can reconcile the experimental results nicely.

### The model reproduces several over-expression phenotypes for NSC’s in CNS

Experimentally it has been shown that over-expression of *Id1* gene inhibited neurogenesis^1,2,40,41^ and model simulation at low doses of BMP2 (where neurogenesis is expected for WT NSC’s (CNS), Fig. 2b) under overexpressed *Id1* condition leads to gliogenesis with a very low expression level of Mash1 protein (Fig. 3g). It is well established experimentally that over-expressing *Mash1* and *Hes6* genes separately will cause increased level of neurogenesis in both the situations^2,6^. The model simulations for *Mash1* (Supplementary Fig. 3h) and *Hes6* (Supplementary Fig. 3i) over-expressions are in excellent qualitative agreement with experimental phenotypes. Model further accurately predicts that simultaneous over-expression of *Ngn* and *Hes6* genes (Fig. 3h) will increase the level of neurogenesis as observed in the experiments^6^. Model simulation makes an interesting prediction for the over-expression phenotype where both *Ngn* and *Hes5* genes are simultaneously over-expressed and it leads to inhibition of neurogenesis for low (Fig. 3i) as well as high (Supplementary Fig. 3j) doses of BMP2. These results perfectly agree with the experimental observations made by Fior et al.^6^. Over-expression of *E47* gene increases the Mash1 stability and expression levels in experiments^2^. Our model simulation can validate this observation for both high (Supplementary Fig. 3k, right panel) and low (Supplementary Fig. 3k, left panel) BMP2 doses. Further in accordance with experimental findings^6,42,43^ we observed that over-expressing negative regulators of neurogenesis such as *Hes1* and *Hes5* genes individually (Supplementary Fig. 3l-m) or simultaneously (Supplementary Fig. 3n) in model simulation will cause inhibition of neurogenesis.

### The model predicts the possible routes to reconcile the developmental features of WT NSC’s in PNS

The proposed model seems quite efficient to qualitatively describe the developmental fate choice of NSC’s related to CNS under different doses of BMP2 for varied biological conditions. Still the question remains, whether the same model can qualitatively delineate the distinctive features those are observed for the WT NSC’s obtained from PNS for different doses of BMP2 (i.e., low dose of BMP2 preferentially leads to gliogenesis and higher dose of BMP2 leads to neurogenesis) or not? We begin to address this question with a hypothesis elucidated in Fig. 4a by a schematic bifurcation diagram of the total Mash1 protein as a function of BMP2 dose, where solid red line corresponds to the CNS like situation. From Fig. 4a, intuitively one can perceive that the positions of the saddle nodes SN_1_ and SN_4_ critically control the dynamics of the overall system as a function of BMP2 dose. If one manages to tune the dynamics of the overall system in such a way that the total system can be represented by a slightly altered bifurcation diagram shown as dashed red line in Fig. 4a, the system can manifest the PNS like behavior. Under such a situation, at low level of BMP2 (2 s.u.) we can also have gliogenesis (with low level of total Mash1 and high level of Hes5 protein) and higher level of BMP2 (20 s.u.) will cause neurogenesis (with higher level of total Mash1 and low level Hes5 protein). Now the question is, how to tune the dynamics of the system to have this kind of effect?

**Figure 4.**
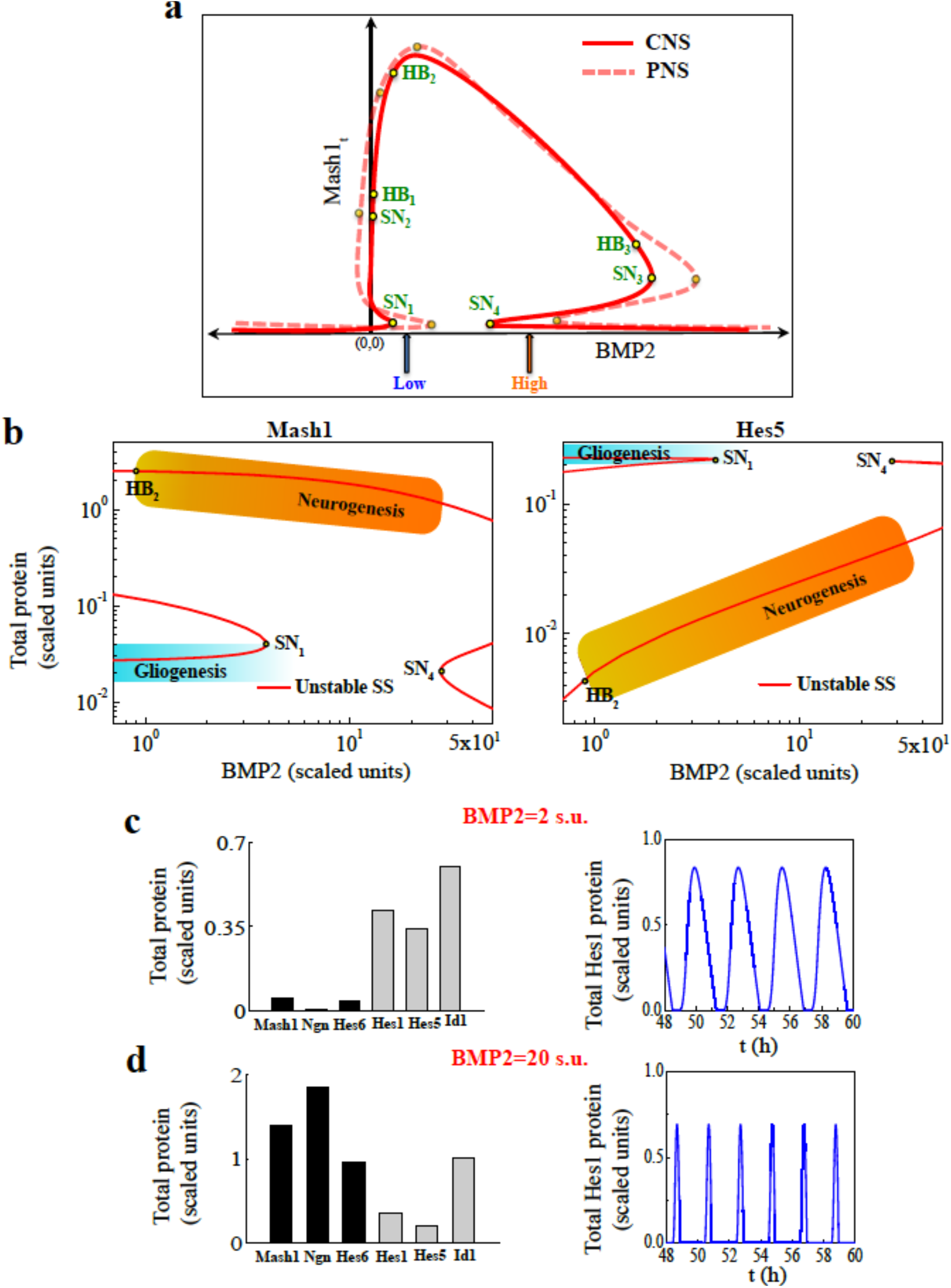
Model predicts the possibility of BMP2 driven developmental lineage commitment of WT NSC’s in PNS. **(a)** A hypothesis is put forward by using the schematic bifurcation diagrams of total Mash1 protein dynamics in CNS (Red solid line) and PNS (Red dashed line) as a function of BMP2 that if the positions of the saddle nodes SN_1_ and SN_4_ (as depicted in red solid line) can somehow be altered simultaneously further towards the higher BMP2 levels (as shown in red dashed line), the system can show a PNS like behavior under the same pre defined low (blue arrow) and high (orange arrow) BMP2 doses. **(b)** Bifurcation diagrams of total Mash1 (left panel) and Hes5 (right panel) proteins in PNS are plotted as a function of BMP2 (both the axes are shown in log scale). Saddle nodes SN_1_ and SN_4_ appear at ∼ 3.91 s.u. and ∼ 27.79 s.u. of BMP2 respectively. Now increase in BMP2 induces the developmental cell fate to change from gliogenic state (low Mash1 and high Hes5 expressions, shown by the graded blue region) to neurogenic state (high Mash1 and low Hes5 expressions, shown by the graded orange region). **(c)** Expression levels of the proneural (Mash1, Ngn and Hes6), gliogenic (Hes5 and Id1) and Hes1 proteins at BMP2=2 s.u. show clear indication of gliogenic fate commitment (left panel) and Hes1 protein concentration shows ∼2 h oscillation period (right panel). **(d)** Expression levels of the proneural (Mash1, Ngn and Hes6), gliogenic (Hes5 and Id1) and Hes1 proteins at BMP2=20 s.u. evidently show neuronal fate commitment (left panel) and Hes1 protein concentration shows ∼2 h oscillation period (right panel). *k*_bmp2_=100 min^-1^ and *k*_bmp22_=3e-02 min^-1^ are used to get PNS like features (b-d). Other parameters are same as depicted in SI Text.

Since, in this study we are more interested to decipher the role of BMP2 on developmental fate determination, we first focused on the BMP2 related parameters *k*_bmp2_, *k*_bmp21_ and *k*_bmp22_ in the model. Our proposed model predicts that by reducing both *k*_bmp2_ (∼20 times) and *k*_bmp22_ (∼400 times) from the values used to describe the WT type case of the CNS, we can have the bifurcation diagrams (Fig. 4b left panel, Mash1 and right panel, Hes5) describing the WT situation of the NSC’s obtained from PNS under different doses of BMP2 as envisioned earlier in Fig. 4a (red dotted line). The corresponding expression levels of proneural and gliogenic proteins indicate the gliogenic fate commitment at 2 s.u. (low dose) of BMP2 (Fig. 4c, left panel) and neuronal fate at 20 s.u. (high dose) of BMP2 (Fig. 4d, left panel). The Hes1 protein concentration still oscillates for both low (Fig. 4c, right panel) and high (Fig. 4d, right panel) dose of BMP2. The model can make further important predictions on different mutation phenotypes, over-expression phenotypes and phenotypes under other important biological conditions for neural precursor cells in PNS (see *SI Appendix* and Supplementary Fig. 4 for details).

### Sensitivity analysis of the model parameters

Interestingly, changing *k*_bmp2_, *k*_bmp21_ and *k*_bmp22_ individually (or changing them in any other binary combination, except the one described above) will not give the desired PNS like feature. What is the reason behind such an observation? Can we have a situation where by only changing any one of the parameters related to the system, we can alter CNS like dynamics to a PNS like one? Can we predict more such possibilities to tune the dynamics? To answer these intriguing questions, we performed a systematic sensitivity analysis (Supplementary Figs. 5 and 6) of all the parameters present in the model (SI Text) by taking the position of the saddle nodes SN_1_ and SN_4_ (Fig. 2a, Supplementary Fig. 2a) separately as sensitivity parameters (For details see methods section). Sensitivity analysis revealed that the proposed model has two distinct kinds of parameters. Increasing any parameter individually will cause the dynamical movement of both the saddle nodes (i.e., SN_1_ and SN_4_) in the opposite direction. Essentially, increasing one kind of parameters individually will take both the saddle nodes (SN_1_ and SN_4_) away from each other. The saddle nodes will eventually approach each other due to individual increment made in the other kind of parameters. Unfortunately, neither of these situations will help us to realize a PNS like feature. This immediately indicates the fact that by only changing one of the parameters of this system will not allow to transform the CNS like system to a PNS like situation.

We have already shown that by varying two parameters (*k*_bmp2_ and *k*_bmp22_) we can have a PNS like behavior (Fig. 4b) and by thoroughly analyzing the sensitivity plots (Supplementary Figs. 5 and 6) we can even show such features for few other pairs of parameters as well (for details see *SI Appendix* and Supplementary Fig. 7). To figure out the rational behind such a choice of parameters, in Fig. 5a, we have compared the sensitivities of the two saddle nodes (SN_1_ and SN_4_) for the two concerned parameters *k*_bmp2_ and *k*_bmp22_. From Fig. 5a it is clear that the saddle node SN_1_ is more sensitive with respect to parameter *k*_bmp2_ than the saddle node SN_4_, whereas the situations are just the opposite when we compare the sensitivities of SN_1_ and SN_4_ as a function of *k*_bmp22_. This gives an immediate hint that tuning these two parameters can modify the dynamics of the system and we eventually get the PNS like features by reducing both of them as described earlier.

**Figure 5.**
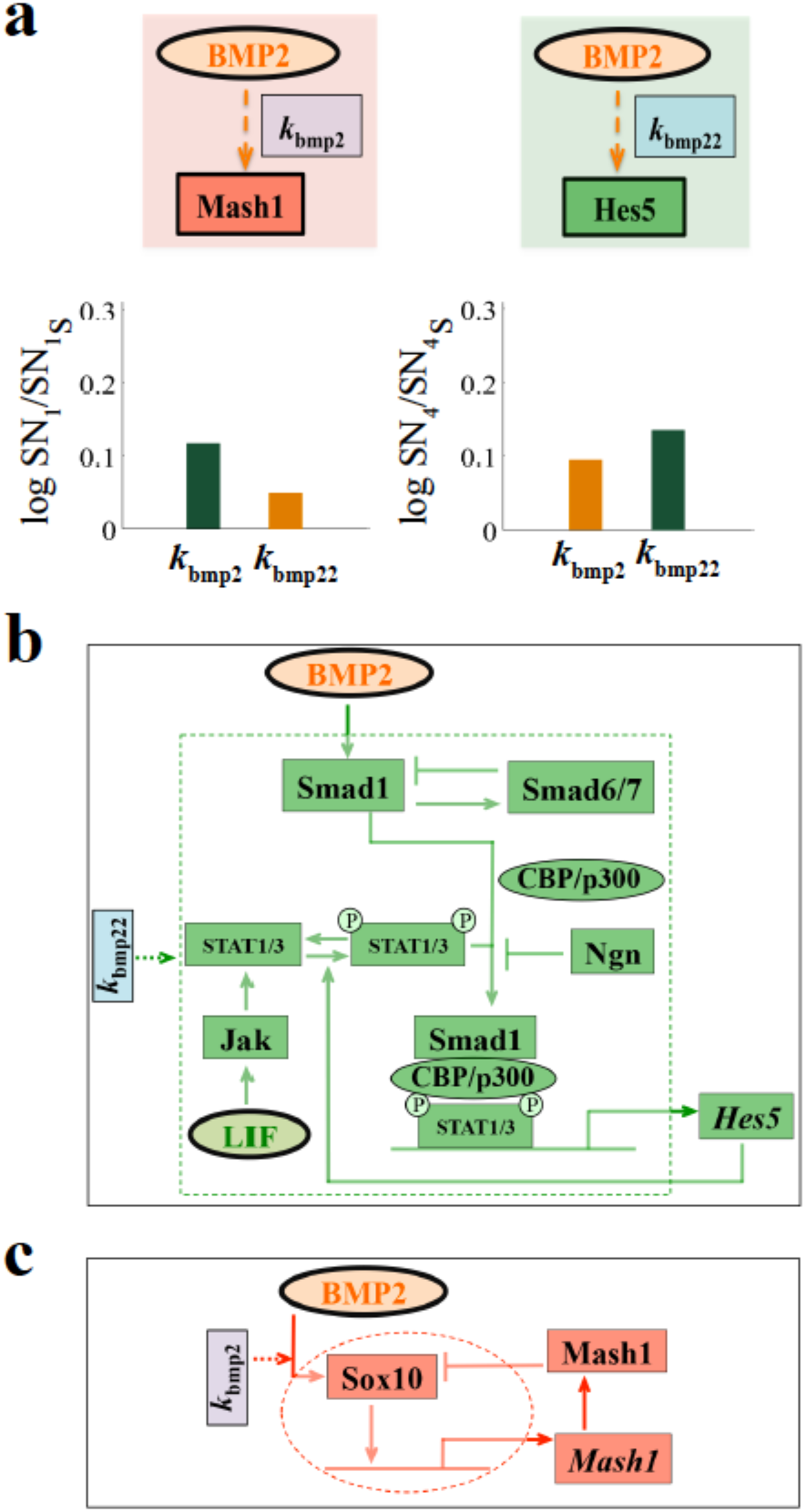
Origin of the differential behavior of NSC’s in PNS in comparison to CNS. **(a)** Upper left and right panels define *k*_bmp2_ and *k*_bmp22_ as the transcriptional activation rates of Mash1 and Hes5 by BMP2. Lower left and right panels show sensitivities of *k*_bmp2_ and *k*_bmp22_ towards SN_1_ and SN_4_ (in Fig. 2a) for WT NSC’s in CNS. *k*_bmp2_ is more sensitive towards SN_1_ and *k*_bmp22_ is more sensitive towards SN_4_. Orange bar signifies movement of the saddle node towards higher BMP2 and dark green bar signifies movement of the saddle node towards lower BMP2 than the WT CNS case. Both the parameters are increased individually by an amount of 20% of the actual model parameters used in Fig. 2a (SI Text) keeping all other parameters constant. **(b)** The detailed regulatory network responsible for BMP2 driven Hes5 transcriptional activation that is modeled in terms of the parameter *k*_bmp22_ in our model. **(c)** The detailed regulatory network responsible for BMP2 driven Mash1 transcriptional activation that is modeled in terms of the parameter *k*_bmp2_ in our model.

### Biological relevance of the model predictions

The important question is, do the existing experimental literature support such kind of prediction? If not, how one should proceed to verify such a prediction experimentally? The biological relevance of reducing *k*_bmp2_ and *k*_bmp22_ getting a PNS like behavior is extremely fascinating and complex. Although in the proposed model we have assumed BMP2 transcriptionally activates both the *Hes5* and *Mash1* genes at a rate represented by *k*_bmp22_ and *k*_bmp2_ respectively, the underlying molecular biological details are quite complex as shown in Figs. 5b-c. Fig. 5b depicts that BMP2 directly activates Smad1, which in turn makes a complex with CBP/p300^16,44^. Phosphorylated form of STAT1/3 can further form a trimeric complex with the Smad1-CBP/p300 complex and the trimeric complex can now act as the transcriptional activator of *Hes5* gene^16^. This linear activation process of *Hes5* gene is controlled at different points in the network. Smad1 activates Smad6/7 that negatively regulates Smad1^44^. Ngn can bind with Smad1-CBP/p300 complex to activate its own transcription and reduces the amount of trimeric complex (between Phosphorylated STAT1/3 and Smad1-CBP/p300) formation^16^. To make things more complex, Hes5 can further activate the LIF mediated formation of phosphorylated STAT1/3 to boost its own transcription^17,44,45^. In our model we represented this underlying network by *k*_bmp22_. Similarly, Fig. 5c shows that BMP2 induces the activation of HMG-box factor Sox10, which is necessary for transient activation of *Mash1* gene^18,46^. Further, it had been shown experimentally that the Mash1 protein either directly or indirectly regulates the expression level of Sox10 protein negatively^18,46^. Again in our model we capture all these regulations shown by the dotted red circle in Fig. 5c by the *k*_bmp2_ parameter. Thus, the prediction that we need to reduce both *k*_bmp2_ and *k*_bmp22_ to get a PNS like feature, tacitly means that the expression level of one or some of the proteins involved in these underlying networks represented by *k*_bmp22_ (Fig. 5b) and *k*_bmp2_ (Fig. 5c) are changing in such a way that dynamically the CNS like system is becoming more PNS like.

In accordance with the observation made by Kim et al.^46^, higher expression level of Sox10 (represented by higher value of *k*_bmp2_ taken in the model for the WT situation in CNS) even in presence of low level of BMP2 can transiently take the Mash1 protein to a high level very quickly (Fig. 2a) and can initially favor the neuronal state. At the same time, higher value of *k*_bmp22_ (which correlates with higher effect of BMP2-Smad signaling) ensured the onset of inhibition of neuronal differentiation by degrading the Mash1 protein with increasing doses of BMP2 in NSC’s in CNS that also complies with the experimental findings in literature^1,16,17,44,45^. This observation is in line with the bifurcation feature of total Mash1 protein shown in Fig. 2a where at lower BMP2 we have neurogenesis (higher Mash1 level) and then at relatively higher values of BMP2 the system prefers the gliogenic fate (low levels of Mash1).

On the contrary, relatively lower values of *k*_bmp2_ (can be correlated with a low expression level of Sox10) and *k*_bmp22_ (can be correlated with a lower effect of BMP2-Smad signaling) will delay the BMP2 driven sudden activation of total Mash1 protein level by pushing the saddle node SN_1_ for relatively higher values of BMP2 and keep the system in the gliogenic state (low Mash1 level) as shown in Fig. 4b (left panel).

Thus, increasing BMP2 doses will eventually drive the system towards neurogenesis (Fig. 4b) for neural crest stem cells obtained from developing PNS. Under these circumstances it will look like that BMP2-Smad signaling is initiating the neuronal differentiation via induction of Mash1 that clearly supports the experimental observation made for developing neural crest stem cells^1,3–5,18,46,47^. The model predicted switch like transition from gliogenic to neuronal fate (higher Mash1 level) as a function of BMP2 doses (Fig. 4b) in neural crest stem cells corroborates with the fact that experiments performed with this cell type showed sudden increase in proportions of neuronal population with increasing doses of BMP2^4^.

## Discussion

Despite some inherent differences that exist between NSC’s obtained from CNS and PNS, it is believed that in these NSC’s of slightly different origin, similar kind of mechanism regulates the cell fate decision-making process during development^3,30,47^. At the same time, experiments in the literature demonstrated that BMP2 differentially regulates the process of neuronal differentiation of NSC’s in central and peripheral nervous systems^1–5^. In this article, we took a mathematical modeling approach to decipher the underlying dynamics that leads to such a unique BMP2 driven cell fate decision-making process operative in NSC’s of central as well as peripheral nervous systems.

To begin with, we took a modular approach (Fig. 1, Supplementary Fig. 1) to build our complete mathematical model (See *SI Appendix*) where we analyzed each module with the help of bifurcation analysis to obtain certain experimentally observed features^1–5^ and later integrated all the modules one by one to obtain the whole model. First, we showed that the mathematical model could describe the WT behavior of the NSC’s in CNS (Fig. 2). It is important to note that our model not only gets the steady state feature corresponding to a developmental fate correct (for example, high Mash1 total protein level at low BMP2 corresponds neurogenesis, Fig. 2a) but it also accounts for the experimentally observed oscillations in Hes1 protein with ∼ 2 hour time period (Figs. 2b-c)^27,32^. Secondly, the model developed for CNS successfully reproduced several experimental observations (Fig. 3, Supplementary Fig. 3 and Table - 1) related to single and double mutant phenotypes as well as over-expression phenotypes for NSC related to CNS qualitatively^2,6,7,16,20,34-43^. Moreover, the model can demonstrate even more subtle experimental observations such as *Hes1* mRNA and protein oscillatory period as well as the amplitude of oscillations of Hes1 almost remain unaltered in both neuronal as well as in the gliogenic state corresponding to low and high BMP2 doses (Figs. 2b-c, 3a, Supplementary Fig. 3a)^1,27,32^.

At this juncture, we envisaged (Fig. 4a) that the presence of two interconnected bi-unstable switches in the steady state dynamics for the important differentiation regulatory proteins (Mash1, Hes5 etc.) as a function BMP2 dose might be the crucial determining factor of the different behavioral pattern that we observe as the doses of BMP2 increases in the neural stem cell medium for central and peripheral nervous systems. With the help of systematic sensitivity analysis on the saddle nodes SN_1_ and SN_4_, we have evidently shown that by changing only one parameter related to the system, it is not possible to convert the dynamics of the system from a CNS to PNS like. The sensitivity analysis also suggests that by carefully tuning a pair of parameters present in the model, it is possible to obtain the required PNS like feature (Fig. 4b, Supplementary Fig. 7).

What it meant biologically was not that straight forward to rationalize. It turned out that the way we modeled the effect of BMP2 on *Mash1* and *Hes5* transcription by using parameters *k*_bmp2_ and *k*_bmp22_ respectively are too simplified (Figs. 5b-c). On one hand, reducing the *k*_bmp2_ can be correlated to reduced expression level of Sox10 protein or some effect that changes the negative regulation of Mash1 protein on Sox10 (Fig. 5c). Whereas, it is hard to correlate the reduction in *k*_bmp22_ directly to any one protein involved in the interaction network shown in Fig. 5b but there are evidences that changing the expression levels of Smad proteins and inducing LIF mediated STAT signaling can alter the lineage specific differentiation regulation for central as well as peripheral nervous systems^16,17,44,45,47^. Thus, the marked differences that we observe in BMP2 driven dynamical response of NSC’s of CNS and PNS can be caused by several routes; (i) the expression levels of different proteins involved in Figs. 5b-c may get altered due to specific epigenetic regulations (ii) Extent of positive and negative regulations operative in the biological network shown in Figs. 5b-c may change to alter the dynamics or (iii) Additional signaling pathways like LIF mediated signaling pathways may work together with BMP2 signaling to differentially regulate cell fate decision making process in CNS and PNS.

The proposed mathematical model is one of the very first attempts to successfully demonstrate BMP2 driven developmental dynamics of NSC’s in CNS, which also accounts for a number of experimentally observed mutant and over-expression phenotypes. Through systematic bifurcation and sensitivity analysis of the proposed model, we discussed few plausible routes that the system might adopt to manifest the differential developmental dynamics of neural crest stem cell in PNS. The proposed mechanism that underlies the shift in the BMP2 driven dynamics in stem cells of central and peripheral nervous systems and other model predictions seem to evoke interesting and challenging questions, which requires further research both in terms of experiments and theory. For example, if bi-unstable switches were governing the differentiation dynamics of NSC’s in both CNS and PNS (as suggested by our proposed mechanism) then one would expect that intrinsic fluctuations present in the system (for example, due to low copy number of mRNA’s) might generate mixed population of neuron and glial cells under intermediate doses of BMP2. Indeed experiments in literature supports this observation for neural crest stem cells in PNS^3–5^.

We believe more experiments in this direction will be crucial to understand the dynamics in a better way and will allow us to explore possible therapeutic strategies to minimize unwanted gliogenesis that results as the aftereffect of nerve injury and neural stem cell implantation. More importantly it will throw new challenges to the proposed mathematical model that has shown a lot of potential to generate exciting questions both in terms of experiments and theory. We hope that in future the model will help us to progress systematically to find an efficient route for neuronal regeneration from NSC’s.

## Methods

### Deterministic simulations

The complete gene regulatory network (Fig. 1, Supplementary Fig. 1) was expressed in terms of 46 ordinary differential equations (SI Text). Module-1 and extended model (*SI Appendix*) were expressed in terms of 13 and 34 ordinary differential equations respectively. The deterministic models (*SI Appendix* and SI Text) were encoded as .ode files and the deterministic simulations were performed using freely available software XPP-AUT. The bifurcation diagrams were drawn by the AUTO facility available in XPP-AUT software. The bifurcation diagrams and time profiles were drawn in OriginLab and MATLAB using the data points generated by XPPAUT.

In the sensitivity analysis section parameters were increased individually at an amount of 20% of the model parameters (SI Text) keeping all other parameters constant and the bifurcation diagrams were drawn by the AUTO facility available in XPP-AUT software. Sensitivity analysis of the model parameters (SI Text) was done on the saddle nodes SN_1_ and SN_4_. The bar diagrams were drawn in MATLAB. Orange bar signifies movement of the saddle nodes towards higher BMP2 and dark green bar signifies movement of the saddle nodes towards lower BMP2 than the WT CNS case.

## Acknowledgements

Thanks are due to IRCC, IIT Bombay (13IRTAPSG005) for a fellowship to (DS). This work is supported by the funding agencies IRCC, IIT Bombay (13IRCCSG008), DST grant (EMR/2014/000500) and DBT grant (BT/PR11932/BRB/10/1315/2014).

## Author Contribution

DS and SK designed the research problem, DS developed the mathematical model together with SK, DS and SK analyzed the modeling outputs and predictions, DS and SK wrote the paper together.

